# Reef environments shape microbial partners in a highly connected coral population

**DOI:** 10.1101/2020.10.27.357095

**Authors:** NG Kriefall, MR Kanke, GV Aglyamova, SW Davies

## Abstract

Corals from more thermally variable environments often fare better under thermal stress compared to those from less thermally variable environments, an important finding given that ocean warming threatens corals worldwide. Evidence is mounting that thermal tolerance can be attributed to the coral itself, as well as microbial communities present within the holobiont (coral host and its associated microorganisms). However, few studies have characterized how thermally variable environments structure multiple holobiont members *in situ*. Here, using 2b-RAD sequencing of the coral and metabarcoding of algal (ITS2) and bacterial (16S) communities, we show evidence that reef zones (locales differing in proximity to shore, physical characteristics, and environmental variability) structure algal and bacterial communities at different scales within a highly connected coral population (*Acropora hyacinthus*) in French Polynesia. Fore reef (more stable) algal communities were on average more diverse than the back reef (more variable), suggesting that variability constrains algal diversity. In contrast, microbial communities were structured on smaller scales with site-specific indicator species and enriched functions across reef zones. Our results illuminate how associations with unique microbial communities can depend on spatial scale across highly dispersive coral populations, which may have fitness consequences in thermally divergent regions and rapidly changing oceans.

## INTRODUCTION

Anthropogenic climate change is causing widespread concerns across ecosystems. Equatorial marine ectotherms, already living near their thermal limits, are especially vulnerable (1). As a vivid example, tropical coral reefs are swiftly succumbing to the consequences of ocean warming (2,3). Rising temperatures induce coral bleaching, a breakdown of the symbiosis between corals and algae (Symbiodiniaceae) (2,3). If thermal stress is not relieved, bleaching and subsequent lack of algal-derived nutrients can lead to coral death (2,3). Widespread efforts are underway to identify natural climate refugia as conservation and management targets (4). Notably, areas of high frequency thermal variability are associated with coral bleaching mitigation (2,3), however, the mechanisms underlying this phenomenon remain unclear.

The potential for adaptation and/or acclimatization to environmental variation spans multiple members of the coral holobiont (unit encompassing coral hosts and their microorganisms) (5,6). Previous work has revealed genes associated with heat tolerance in corals from contrasting thermal environments, and that this heat tolerance can be heritable (7,8). In addition, association with certain Symbiodiniaceae has been well-documented to confer bleaching resistance (9,10). Most recently, insights into the coral’s microbiome have revealed putative functions that mitigate the effects of thermal stress, such as nutrient cycling and immunity (11,12). Indeed, inoculating *Pocillopora damicornis* with a consortium of potentially beneficial bacteria prevented bleaching during both heat stress and pathogen challenge (13). Clearly, to better understand coral resilience across environments, a holobiont perspective must be taken.

Reef zones provide an effective natural system to study the coral holobiont under divergent conditions, as strong environmental variation can be found even within 1 km (6,14). Indeed, corals from more variable reef zones can exhibit increased thermal tolerance when compared to corals from more stable reef zones (2,15,16), but see (17). Significant genetic differentiation across reef zones, as well as distinct algal and bacterial communities, has been observed in brooding coral *Pocillopora damicornis* (6), highlighting the potential for multiple holobiont members to be structured across short distances. However, a holistic view of how multiple holobiont members vary across reef zones in a broadcast spawning coral, where genetic differentiation across smaller scales is much less likely (18), remains less explored. In these cases where life histories imply that corals will exhibit longer range migration, associations with unique microbial communities might serve as a mechanism for corals to acclimatize to novel environments.

*Acropora hyacinthus* is a dominant Indo-Pacific broadcast spawning coral that thrives across reef zones (19). These zones differ in temperature, light, and nutrient levels, where the back reef (close to shore) typically experiences greater variability when compared to the fore reef (close to open ocean) (6,20). Here, we leverage reef zones in French Polynesia to ask how each member of the *A. hyacinthus* holobiont is structured across fore reef and back reef environments. Specifically we quantify: (1) coral host genetic structure and associated loci under selection, (2) algal community composition and diversity, and (3) bacterial community composition, diversity, and functional profiles. Together, these data illuminate how these microbial members of the coral holobiont vary across distinct environments and provide potential mechanisms by which those from more variable environments may persist under climate change.

## MATERIALS & METHODS

Detailed materials and methods can be found in Supporting information.

### Coral sampling and site characterization

In 2013, coral branch tips were collected from one fore reef (F) and one back reef (B) zone at each of three sites in French Polynesia: Tahiti (TNW), Mo’orea NW (MNW), and Mo’orea SE (MSE) (Figure 1A; Table S1; CITES export permit #FP1398700064-E). Samples were preserved in 96% ethanol and maintained at −20°C until processing. When possible, each colony was photographed (no MSE-F photographs due to inclement weather) and surface area was estimated using ImageJ (21) (*e*.*g*. Figure 1C). Colony surface area (cm^2^) data were transformed (Yeo-Johnson) and a one-way ANOVA tested for size differences between reef zones.

**Figure 1.**
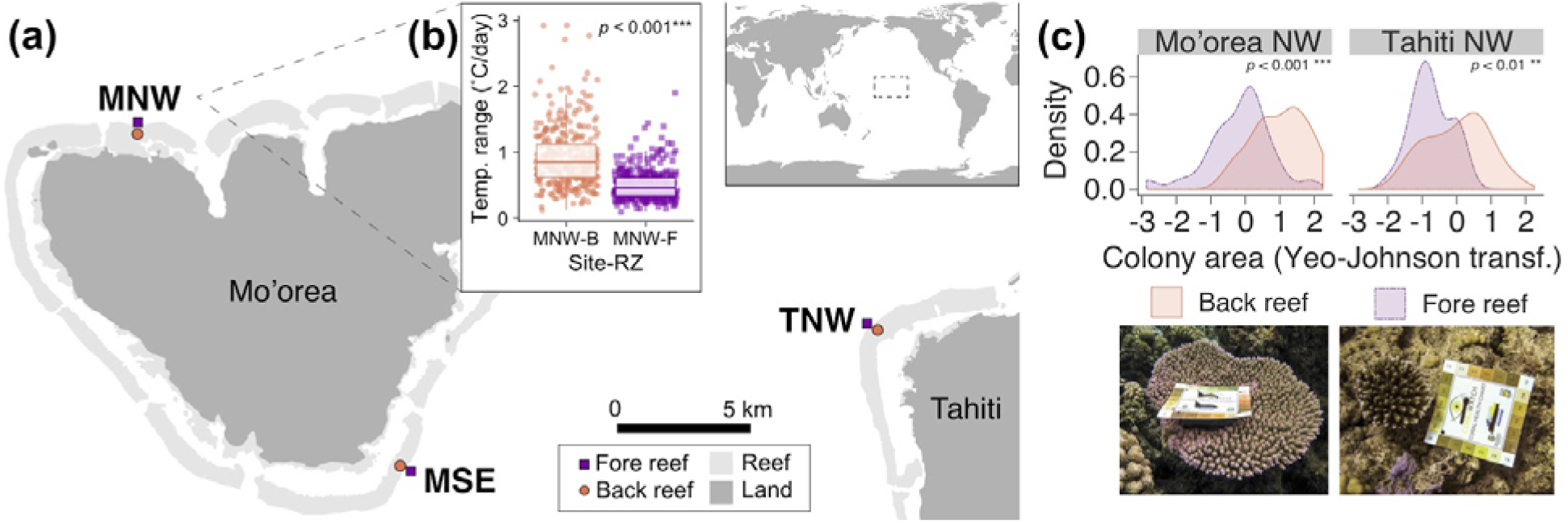
**(a)** Map of French Polynesia study sites: Mo’orea NW (MNW), Mo’orea SE (MSE), and Tahiti NW (TNW) (paired back reef and fore reef zones). **(b)** Daily temperature (°C) ranges at MNW back reef (B) and fore reef (F) from June 2013 – 2014, which overlaps with sampling period (07-08/2013). Data from Service D’Observation CORAIL. **(c)** Density plots of *Acropora hyacinthus* colony surface area (cm2; Yeo-Johnson-transformed) sampled from fore reef and back reef zones in MNW and TNW (no photographic data available for MSE) with representative photographs from back reef (left) and fore reef (right).

Temperature data from two SBE 56 Temperature Sensors (Sea-Bird Scientific) labeled ‘B1’ (MNW-B) and ‘P3’ (MNW-F) sites were provided by Service d’Observation (SO) CORAIL through CRIOBE. Hourly data from June 1^st^, 2013 through June 1^st^, 2014 were extracted. Mean daily temperature and ranges were compared between reef zones using Mann-Whitney *U* tests.

### Host 2b-RAD genotyping

Holobiont tissue was isolated following (22). DNA concentrations were standardized to 25 ng/µl for 2b-RAD genotyping. 2b-RAD library preparations were modified from (23) following https://github.com/z0on/2bRAD_denovo/. Briefly, DNA (100 ng) was digested with *BcgI* (New England Biolabs) and fragments were multiplexed using custom barcoded ligation adapters. Samples were pooled and sequenced on three lanes of Illumina HiSeq 2500 (76 samples; Table S2) or one lane of Illumina HiSeq 4000 (60 samples; Table S2). Ten of 136 samples were technical replicates (Table S2). Raw and processed read count data are available in Table S2.

All bioinformatic analyses are available at https://github.com/nicfall/moorea_holobiont. Sequencing data were demultiplexed, de-duplicated when appropriate (see Table S2), and filtered using FASTX-Toolkit (24). Filtered reads were mapped to the *Acropora millepora* genome (25) using *Bowtie2* (26). Analysis of Next Generation Sequencing Data (*ANGSD*) (27) calculated genotype likelihoods using the following filters: only biallelic calls, only one best hit per read, minimum mapping quality score of 20, minimum base quality score of 25, maximum strand bias *p*-value of 10^-5^, maximum heterozygosity bias *p*-value of 10^-5^, a minor allele frequency of 0.1, a minimum depth per site of 8 reads, and sites were required to be present in at least 80% of samples. Pairwise identity-by-state (IBS) matrices were calculated and results plotted as a distance dendrogram (Figure S1). Technical replicates and clones were removed, leaving only one genotype representative. Samples with an average per-site depth of <7 reads were removed to improve genotype likelihood confidence (*n*=10 removed; Table S2). *PCAngsd* (28) generated a covariance matrix for principal components analysis. Significance of multivariate dispersion and location were assessed with *vegan* functions *betadisper* and *adonis*, respectively (29). The *ANGSD* VCF file was converted into BED format using *PLINK* (30) and run through *ADMIXTURE* (31) (K 1-5) where the lowest cross-validation error determined the optimal ‘K’. *hierfstat* (32) calculated global and pairwise F_ST_ between all sites and reef zones. *BayeScan* (33) and *OutFLANK* (34) were used to identify single nucleotide polymorphism (SNP) F_ST_ outliers.

### Symbiodiniaceae metabarcoding

From a subset (*n*=96; 16/reef zone/site; Table S1) of samples used in host genotyping, ITS2 metabarcoding was performed. ITS2 libraries were generated using a series of PCR amplifications described in (35) and sequenced on Illumina Miseq (paired-end 250bp) at the University of Texas at Austin. ITS2 data were pre-processed in *bbmap* (36) and *cutadapt* (37) removed primer sequences. *DADA2* (38) truncated reads, calculated error rates, de-duplicated reads, inferred sequence variants, merged paired reads, and removed bimeras (38). One sample failed quality filtering (Table S2). 118 non-bimeric amplicon sequence variants (ASVs) were assigned taxonomy using the GeoSymbio database (39) with a minimum bootstrap confidence of 70. LULU curation algorithm (40) combined ASV counts with 99% sequence match. The clustered ASV table was rarefied to 1,994 counts per sample, using *vegan* (29), which retained ∼90% of samples (Table S2). *Phyloseq* (41) calculated three alpha diversity metrics: ASV richness, Shannon index, and inverse Simpson’s index. Mann-Whitney *U* tests assessed significance across reef zones overall and within each site. Linear regressions assessed correlations between alpha diversity metrics and colony size.

*MCMC*.*OTU* (42) trimmed ASVs representing <0.1% of reads or those present in only one sample. A Principal Coordinate Analysis (PCoA) using Bray-Curtis dissimilarity was conducted using *phyloseq* (41). Functions *adonis* and *betadisper* in *vegan* (29) evaluated sample dissimilarity in multivariate space in terms of location and dispersion, respectively (99 permutations), contrasting reef zones and colony size within each site and *pairwise*.*adonis* (from https://github.com/Jtrachsel/funfuns; 99 permutations) contrasted communities between sites. All analyses were repeated without rarefaction, using relative abundances to confirm results. In relative abundance analyses, all but three samples were retained whose counts were at least 2.5 standard deviations lower than the mean (Table S2).

### Bacterial metabarcoding

The same ITS2 sample subset was used to characterize the coral’s microbiome using 16S metabarcoding, except three samples which were substituted due to insufficient material and three samples that failed to amplify (*n*=93, 15-16/site; Table S1, Table S2). The V4/V5 region was amplified following (43). Pooled libraries were sequenced on an Illumina Miseq (paired-end 250 bp) at North Carolina State University’s Genomic Sciences Library.

Pre-processing and *dada2* analyses followed ITS2 protocols described above. 16S taxonomy was assigned using the Silva v132 dataset (44). Using *phyloseq* (41), ASVs assigning to family “Mitochondria”, order “Chloroplast”, or those failing to assign to kingdom “Bacteria” were removed. Using *vegan* (29), the ASV table was rarefied to 12,000 reads, which retained ∼90% of samples (9 samples removed; Table S2). *Phyloseq* (41) calculated three diversity metrics: ASV richness, Shannon index, and inverse of Simpson’s index. ASV richness and inverse Simpson index were log-transformed and a one-way ANOVA with Tukey HSD tests compared diversity metrics across sites and reef zones. *MCMC*.*OTU* (42) then trimmed ASVs representing <0.01% of counts or only present in one sample. Linear regressions assessed correlations between alpha diversity metrics and colony size.

Sample dissimilarity was evaluated for the total bacterial community as described for ITS2. These analyses were then repeated on 16S data separated by *microbiome* (45) into core (present in >70% of samples) and accessory components, and finally repeated using unrarefied, relative ASV abundances to confirm results. *Indicspecies* (46) (999 permutations) identified significant associations between ASVs and reef zones at each site, after multiple test correction (47). *Piphillin* (48) predicted sample metagenomic content using the Kyoto Encyclopedia of Genes and Genomes (KEGG) Database (49) and *DESeq2* (50) identified differentially enriched metagenomic content across reef zones within sites.

## RESULTS

Detailed results can be found in Supporting information.

### Site and coral colony characterization

Daily temperature range was higher in back reef relative to fore reef environments at Mo’orea NW (Figure 1B). In contrast, mean daily temperature did not vary between reef zones. Coral colony area was higher at the back reef than the fore reef at both Mo’orea NW and Tahiti NW (Figure 1C).

### Coral genetic structure and loci under selection

After quality filtering, reads per sample averaged 1.36±0.85 million (mean±SD) (Table S2). Mean mapping efficiency across samples was 81.97±4.03 % (mean±SD) (Table S2). The following samples were removed: 10 technical replicates (Figure S1), 2 incidental clones (Figure S1), and 10 samples having mean read depth <7/site (Table S2). 114 samples passed quality filtering, which resulted in 3,594 SNPs. *ADMIXTURE* (31) determined that the optimal K=1 (Figure S3). Principal component analysis confirmed this pattern, with sample overlap across sites and reef zones (Figure S2). No differences in either multivariate location or dispersion across sites, reef zones, or their interaction were observed (Figure S2). *BayeScan* (33) and *OutFLANK* (34) found no F_ST_ outliers. Pairwise F_ST_ values between sites ranged between 0.011 and 0.018 (Table S3) and global F_ST_ was 0.001.

### Symbiodiniaceae community composition

Average ITS2 counts were 4,681±2,574 (mean±SD) (Table S2). Most samples were dominated by one of four ASVs assigned to ITS2 type *Cladocopium* C3k (mean relative abundance 80.8%). The second most abundant ITS2 type was *Cladocopium* Cspc (mean relative abundance 12.5%). Background symbiont types included two undescribed *Cladocopium* ASVs and one *Symbiodinium* A1 ASV. Within all three sites, fore reef and back reef communities were significantly different (Figure 2A). Data were not significantly dispersed in multivariate space, with the exception of Tahiti NW, where back reef data were more dispersed than fore reef (Figure 2A). Colony size significantly impacted community data at Tahiti NW only. With reef zone data combined by site, algal communities were different between Mo’orea NW and Tahiti NW, and between Mo’orea SE and Tahiti NW, but not between Mo’orea SE and Mo’orea NW (Figure S4B).

**Figure 2.**
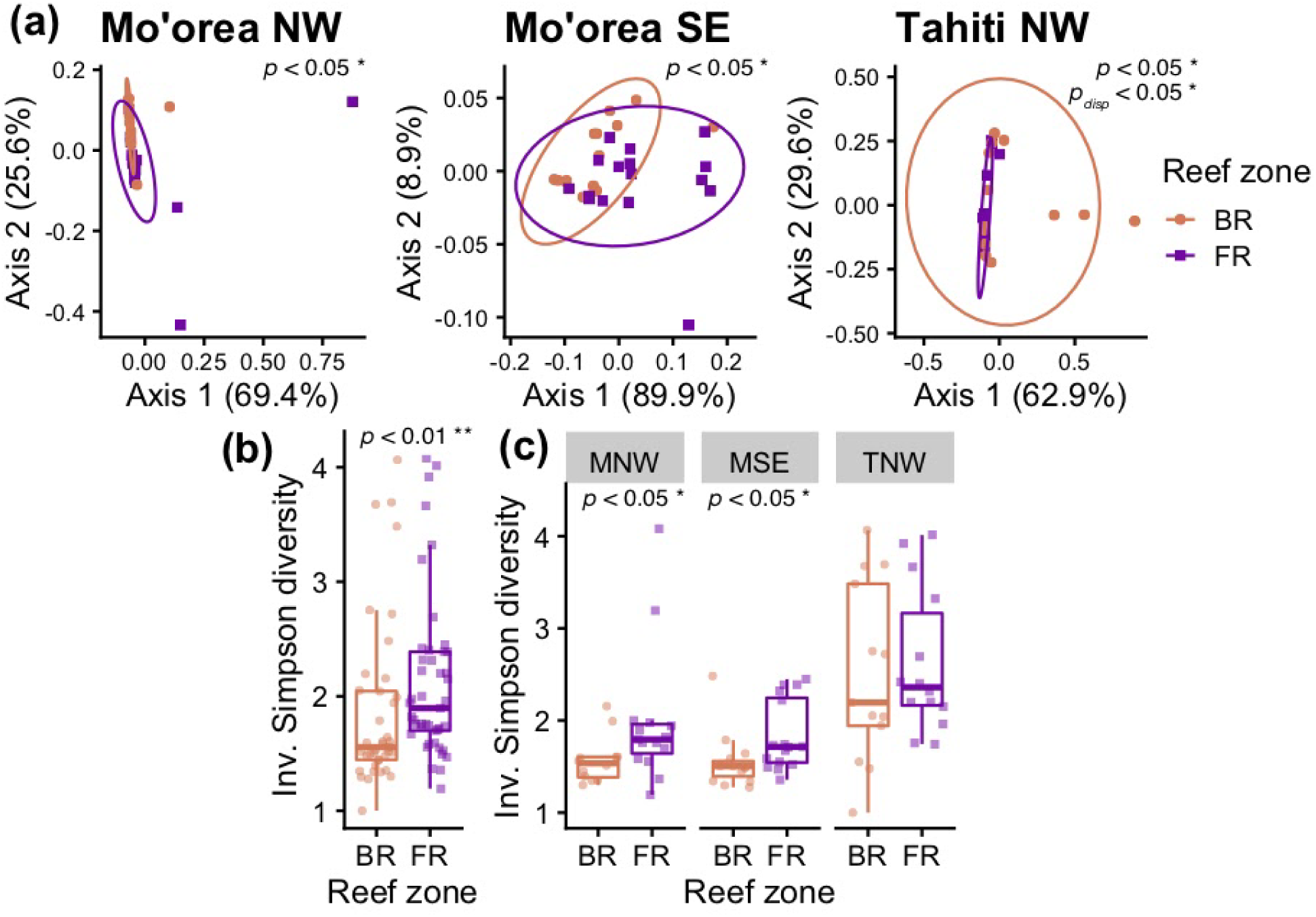
**(a)** Symbiodiniaceae communities in *Acropora hyacinthus* across reef zones (BR: back reef, FR: fore reef) using multivariate ordination plots (PCoA) based on Bray-Curtis dissimilarity. *P-*values indicate a significant comparison of multivariate location, while *pdisp*-values indicate a significant comparison of dispersion. **(b-c)** Inverse of Simpson’s diversity compared across reef zones, **(b)** all sites combined and **(c)** within sites (MNW: Mo’orea NW, MSE: Mo’orea SE, TNW: Tahiti NW.)

Two metrics of alpha diversity indicated higher diversity in fore reef algal symbionts when compared to back reef (Figure 2B). Within each site, there was higher diversity in fore reef samples at Mo’orea NW and Mo’orea SE, but no differences observed between reef zones at Tahiti NW (Figure 2C). There was only a significant difference in ASV richness across reef zones at Mo’orea SE, with higher ASV richness in the fore reef. Colony size did not correlate with any diversity metrics.

### Bacterial community composition

Mean 16S coverage was 27,340±12,999 (mean±SD) prior to rarefaction. Of 207 ASVs, 10 comprised the core microbiome (Table S4). The most abundant member was an ASV belonging to the Phylum Proteobacteria (35.76%). The next two most abundant core ASVs assigned to *Burkholderia-Caballeronia-Paraburkholderia* (23.11%) and *Endozoicomonas* (15%). The remaining core and accessory ASVs averaged <6%.

Shannon diversity, inverse Simpson index, and ASV richness were statistically indistinguishable across reef zones within sites and there was no relationship between diversity metrics and colony size. Multivariate location of 16S communities was significantly different between Mo’orea SE and Mo’orea NW and between Mo’orea SE and Tahiti NW, but not between Mo’orea NW and Tahiti NW (Figure S7).

Within each site, 16S communities were significantly different between the fore reef and back reef at Mo’orea SE and Tahiti NW, but not at Mo’orea NW (Figure 3A). There were no significant differences in multivariate dispersion across reef zones or sites. Patterns did not change when examining the core microbiome (Table S4). However, all reef zone comparisons at all sites were significant when examining the accessory microbiome. There was no relationship between colony size and community composition.

**Figure 3.**
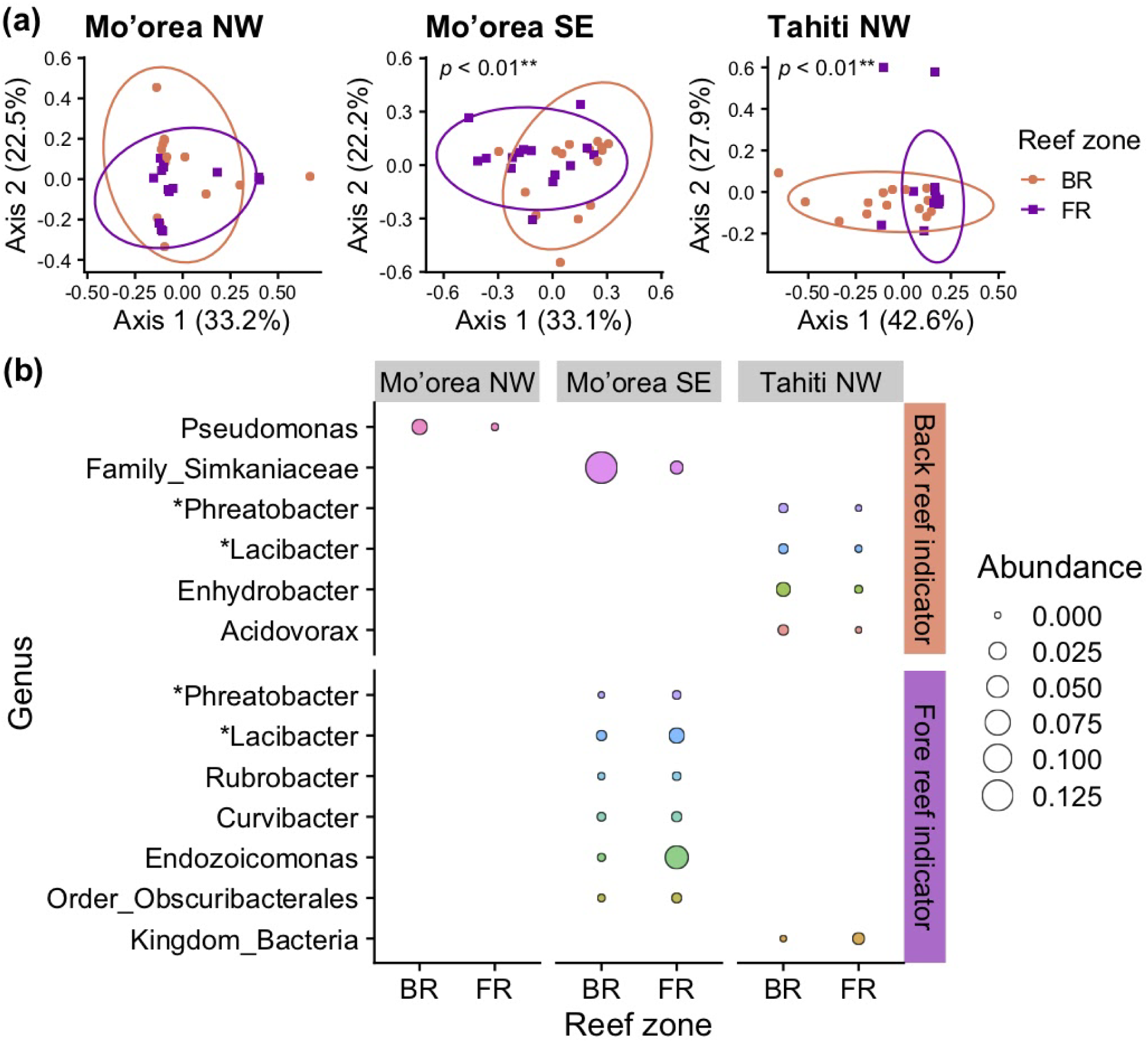
*Acropora hyacinthus* bacterial communities across reef zones (BR: back reef, FR: fore reef) in **(a)** Mo’orea NW **(b)** Mo’orea SE and **(c)** Tahiti NW using multivariate ordination plots (PCoA) based on Bray-Curtis dissimilarity. **(d)** Average relative abundances, scaled with circle size, of indicator bacterial genera. Genera that appeared as indicators of both back reef and fore reef are marked by asterisks (*). (BR: back reef, FR: fore reef.)

Indicator bacterial ASVs for each reef zone were largely unique to each site (Figure 3B). Only one back reef indicator ASV was detected for Mo’orea NW (*Pseudomonas sp*.) and none were detected at the fore reef. One back reef indicator ASV was significant for Mo’orea SE (Family Simkaniaceae) and six for the fore reef (*Phreatobacter sp*., *Lacibacter oligotrophus, Rubrobacter sp*., *Curvibacter sp*., *Endozoicomonas sp*., Order Obscuribacterales). There were four back reef indicator ASVs for Tahiti NW (*Phreatobacter sp*., *Lacibacter oligtrophus, Enhydrobacter aerosaccus, Acidovorax*), and one for the fore reef (Kingdom Bacteria). Interestingly, two ASVs assigned to *Phreatobacter* and *Lacibacter* were back reef indicators at Tahiti NW, but were fore reef indicators at Mo’orea SE (Figure 3B).

Inferred functional profiles showed that functions significantly enriched across reef zones were largely site-specific (Figure 4). Exceptions included: between Mo’orea NW and SE, four functions were conserved and enriched in the back reef (vanillate monooxygenase ferredoxin subunit; L,D-transpeptidase Erfk/SrfK; carboxynorspermidine decarboxylase; LysR family transcriptional regulator, putative pyruvate carboxylase regulator), and one in the fore reef (dipeptidase D); between Mo’orea NW and Tahiti NW, six functions were conserved and enriched in the back reef (aldose sugar dehydrogenase; aminoglycoside 3-N-acetyltransferase I; tetR/AcrR family transcriptional regulator, tetracycline repressor protein; streptomycin 3”-adenylyltransferase; dihydropteroate synthase type 1; small multidrug resistance pump) (Figure 4; Table S5).

**Figure 4.**
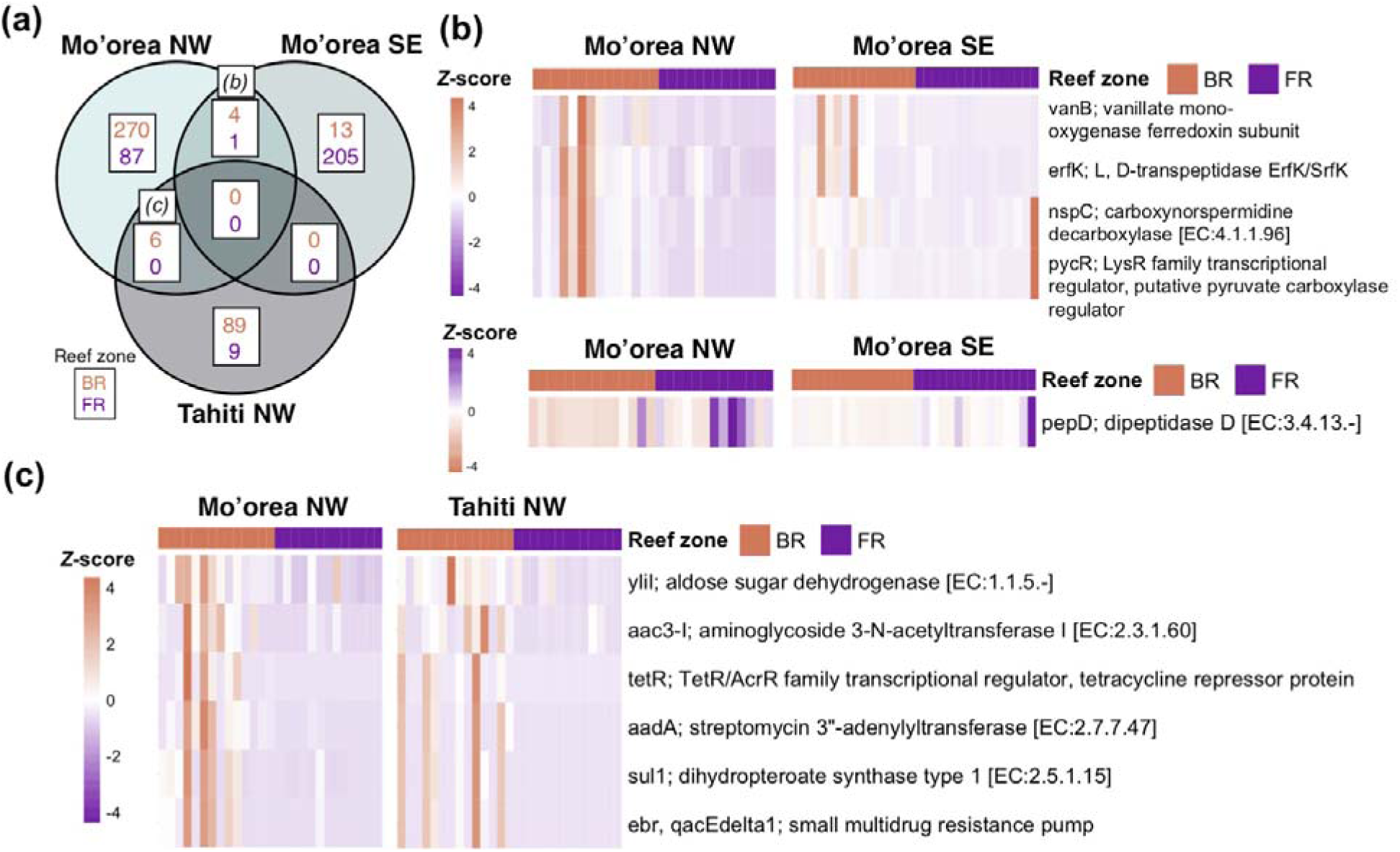
Metagenomic profiles inferred from 16S data. **(a)** Numbers of functional orthologs differentially enriched across reef zones (upper number is back reef, lower number is fore reef) per site. Italicized letters above boxes refer to corresponding panel heat maps. **(b)** Differentially abundant orthologs shared by Mo’orea NW and Mo’orea SE across the back reef (upper heat map) and fore reef (lower heat map). **(c)** Differentially abundant orthologs shared by Mo’orea NW and Tahiti NW across the back reef. (BR: back reef, FR: fore reef.)

## DISCUSSION

### Coral host gene flow is pervasive across reef zones and islands

Studies where dispersive marine organisms display structured populations and outlier loci across surprisingly small distances, likely due to environmental selection or dispersal barriers, are accumulating rapidly (6,7,15). Reef zones can play a notable role in this structuring. Bongaerts et al. (14) and van Oppen et al. (6) found higher differentiation across habitats (reef flat vs. slope; meters apart) than across sites (kilometers apart). In American Samoa, 114 outlier loci were found in *A. hyacinthus* sampled from neighboring back reef pools, even though no overall genetic structure was detected (8). Here, however, we observed that gene flow across habitat types, sites, and even islands was pervasive (Figures S2, S3) and no outlier loci were uncovered, suggesting that this *A. hyacinthus* population experiences high gene flow. This finding supports the growing consensus that the scale of gene flow is largely locale-specific and not easily generalized, even within the same species (51,52).

Other studies examining genetics of Pacific Acroporids have displayed little structure across greater oceanic distances (*i*.*e*. > 100 km) than investigated here (25,53–55). Davies et al. (18) demonstrated strong *A. hyacinthus* population connectivity across thousands of km, which is consistent with our results between islands separated by only ∼15-30 km. However, the genotyping method used here, which re-sequences ∼1% of the genome (23), may have missed putative loci under selection. Future work leveraging whole-genome resequencing or collections from reef habitats with stronger environmental differences, *e*.*g*. (7), may uncover different patterns. Nevertheless, our results indicate that molecular mechanisms beyond the host genome, namely acclimatory changes such as changes in gene expression (16,56,57) or changes in other holobiont components (5,58), are likely responsible for *A. hyacinthus* thriving across divergent these reef zones.

### More diverse Symbiodiniaceae communities in fore reef environments

Symbiodiniaceae communities were co-dominated by *Cladocopium* C3k and Cspc, with background levels of other *Cladocopium* and *Symbiodinium* lineages (Figure S4A), consistent with compositions found in Great Barrier Reef *A. hyacinthus* (59). Differences in relative abundances of ITS2 types (Figures S4, S5) resulted in distinct community compositions across reef zones (Figure 2A). These results are in line with numerous studies where ITS2 communities are structured by reef habitats (6,14,35,60). For instance, environmental variation across reefs in Belize affected the types and abundances of symbionts hosted by *Siderastrea siderea* (35,43). Strikingly, van Oppen et al. (6) found that dominant ITS2 types within *Pocillopora damicornis* were dependent on reef habitat, with C33 dominant on reef slopes and C42 on reef flats. Lastly, coral size is a proxy for age (61), and younger corals often have more diverse symbiont communities than adults (62), however we found no consistent pattern between colony size and ITS2 communities.

We observed higher alpha diversity in Symbiodiniaceae communities hosted by fore reef corals for two of three sites (Figure 2B,C). Most ITS2 types were present in more samples and in higher abundances in the fore reef (Figure S5). It is possible that selective pressures in the back reef, *e*.*g*. increased thermal variability (Figure 1B), may constrain community evenness. As an extreme example of reef environments constraining symbiont types, Bay and Palumbi (2014) found that nearly all *A. hyacinthus* in a highly variable back reef pool were dominated by *Durisdinium* while corals in the moderately variable pool hosted a mix of *Cladocopium* and *Durisdinium*. In contrast, van Oppen et al. (2018) found higher symbiont alpha diversity in *Pocillopora damicornis* living in more variable reef flats, highlighting that diversity patterns may be site or species specific. In *A. millepora*, corals hosting more diverse symbiont communities performed poorer under stress (63), but see (64), an area for follow-up investigations with important implications for coral persistence.

Differences in *A. hyacinthus* Symbiodiniaceae communities may arise from environmental structuring of free-living Symbiodinaceae sources or structuring by the host post-uptake (65). Fine-scale environmental data collection at all sites as well as sampling of Symbiodiniaceae sources would aid in disentangling the contributions of environmental factors to community-level differences. In addition to temperature, nutrients, flow, and water quality have all been suggested to influence Symbiodiniaceae communities across reef environments (35,66,67). Differences in Symbiodiniaceae communities have important implications for the coral, as different species of Symbiodiniaceae can impact host fitness (9,10,68). It is also important to note that ITS2 is multicopy and it is difficult to interpret intra- and intergenomic sequence diversity (69). Therefore, future work should employ more sensitive markers and conduct population-level analyses of *Cladocopium spp*. to better understand algal symbiont structuring across French Polynesia (9,70,71).

### Bacterial taxa and functions are structured on smaller scales

16S total and core bacterial communities were distinct across reef zones at two of three sites (Figure 3A), while accessory bacterial communities were distinct across reef zones at all three sites. These results add to the growing body of literature demonstrating that local reef environments are important drivers of coral microbiome structure (6,43,72,73).Environments differing in thermal variability have been shown to similarly structure bacterial communities, ranging from *A. hyacinthus* in American Samoa (74) to *Siderastrea* spp in Belize (43). However, other factors co-vary with temperature across reef zones (*e*.*g*. flow, water quality, light) and such factors have also been implicated in coral microbiome community assembly (72,75). Potential differences in environmental sources also exist across reefs as distinct bacterial water column communities have been found across reef zones in Mo’orea (20). Further work will be required to disentangle the structuring roles of biotic and abiotics factors for the coral holobiont.

Indicator bacterial taxa for reef zones were specific to each of the three sites and some taxa have been previously suggested to play roles in the coral holobiont (Figure 3B). For instance, members of the genus *Pseudomonas*, an indicator taxon for the Mo’orea NW back reef (Figure 3B), may be a beneficial microorganism in terms of sulfur cycling and antimicrobial activity (12). The genus *Acidovorax*, an indicator taxon in Mo’orea SE back reef (Figure 3B), also has nutrient cycling capabilities through nitrogen-related pathways (76). The genus *Curvibacter* was an indicator in Mo’orea SE fore reef, and this taxon has been previously observed in corals (77,78) and shows high competitive capability in the Cnidarian model *Hydra* potentially due to uptake of host sugars with ABC-transporters (79,80). Finally, genus *Endozoicomonas*, which is found in marine invertebrates and vertebrates worldwide (81), was an indicator taxon for Mo’orea SE fore reef. Within associations with Acroporid corals, *Endozoicomonas* members have shown sensitivity to pH (82,83) and enrichment of KEGG pathways associated with membrane and ABC transporters as well as cell signaling, motility, and secretion (82). Overall, most identities and functional roles for coral-associated bacteria remain unclear and represent a knowledge gap in microbial symbioses generally.

All three sites displayed differentially enriched bacterial functions across their reef zones (Figure 4). Of the enriched functions consistent across reef zones at multiple sites (Figure 4B,C), most shared between Mo’orea NW and Mo’orea SE were involved in metabolism, while half shared between Mo’orea NW and Tahiti NW were involved in antimicrobial resistance (Table S5). Both nutrient cycling and defense against pathogens have been postulated as mechanisms that microbes use to help corals rapidly acclimate to their environment (5,58). Two metabolic functions identified here were related to amino acid metabolism (glutathione and proline/arganine; Table S5) and these are consistent with findings in *A. granulosa* (84). Interestingly, all but one of the functions conserved across multiple sites’ reef zones were enriched in more variable back reef sites. Our results corroborate Ziegler et al. (74), which showed the majority of enriched bacterial functions within *A. hyacinthus* hosts between moderately and highly variable reef pools were observed in the more variable environment, perhaps due to environmental constraints. In addition, nutrient concentrations are typically higher in the back reef around Mo’orea (20), perhaps necessitating the enrichment of functions related to nutrient metabolism. Taken together, specific nutrient exchange and antimicrobial defense represent likely candidates for bacterial functions underlying coral success in the more variable environment; however, much work remains to empirically link bacterial function to host performance (74).

## Conclusion

Understanding the scale of environmental selection for multiple members of the coral holobiont remains critically important as intra-reef variability can determine how corals respond to environmental stressors (2,7). Here, we found evidence of environmentally structured microorganism communities in a highly dispersive coral host. While we did not explore the fitness consequences of hosting these divergent communities, evidence is accumulating that microorganisms can help or hinder the coral host acclimate to rapid environmental change (10,58). This work contributes to this burgeoning area of investigation by providing candidate microbial taxa and functions that may facilitate or impede *A. hyacinthus* acclimatization to divergent conditions. As oceans continue to warm, understanding the distributions and roles of key microbial partners in holobiont resilience may hold the key to effective reef restoration.

## Supporting information

Table S2 sample reads

## ACKNOWLEDGEMENTS

We thank David Lecchini, Véronique Berteaux, and Elina Burns for assistance with fieldwork and permit logistics; Rose Sulentic, Mark Lopez, Nana-Ama Anang, and Lara Laake-Emery for wet lab assistance; the Methods in Ecological Genomic Analysis 2b-RAD class of 2014 for assistance with library preparation; Mikhail Matz for contributions in experimental design, molecular lab space, and bioinformatics; and finally Laura Tsang, Brianna Regan, and John Weldon for help analyzing coral images. Funding was provided to SWD by the Institute des Récifs Coralliens du Pacifique with L’ecole Pratique des Hautes Etudes in addition to a start-up award from Boston University.

## DATA ACCESSIBILITY

Raw 2b-RAD.fastq files are available at NCBI’s SRA (PRJNAXXX). Raw ITS2 and 16S.fastq files are available on NCBI’s SRA (ITS2: PRJNA660421; 16S: PRJNA660779). Processed data files and scripts used in analysis are available at https://github.com/Nicfall/moorea_holobiont.

## AUTHOR CONTRIBUTIONS

SWD conceived of the study. SWD and MRK conducted all fieldwork. NGK, GVA, and SWD conducted all wet lab work. NGK conducted all analyses with help from SWD and MRK. NGK drafted the manuscript and all authors provided comments and approved the final version.

