## Supporting Information for "Reef environments shape microbial partners in a highly connected coral population"

**METHODS**

*Coral sampling*

        	Between July 26^th^ and August 9^th^, 2013, samples were collected from three sites in French Polynesia: Tahiti (TNW), Mo’orea NW (MNW), and Mo’orea SE (MSE), with each site consisting of one fore reef (F) and one back reef (B) zone (see Figure 1A; Table S1). Using snorkeling or SCUBA, small (~2 cm^3^) branch tips from individual coral *Acropora hyacinthus* colonies were collected and colonies were at least 5m apart to avoid sampling clones (see Table S1 for numbers per site). In addition, when possible, each sampled colony was photographed for later species identification confirmation and colony size estimation via surface area analysis (see below). No photographs were obtained at MSE-F due to inclement weather. Sampled branches were preserved in 96% ethanol, kept on ice during transportation to the University of Texas at Austin (under CITES export permit #FP1398700064-E), and maintained at -20˚C until processing.

*Coral colony size assessment*

    	Photographs in the field were taken with a Coral Health Chart (CoralWatch) flush with the colony plate surface (*e.g.* Figure 1C). Photographs were imported into ImageJ (1) and an approximate surface area of the coral was estimated by tracing the total surface outline, with the Coral Health Chart as a size standard to convert the outlined area to cm^2^. Outline measurements of the coral colony and chart were each taken three replicate times and these values were averaged to produce one size measurement per colony. Using R (2), coral colony surface area (cm^2^) data were transformed using the Yeo-Johnson method to meet assumptions of normality, as determined by package *bestNormalize* (3). A one-way ANOVA tested for size differences between reef zones at Mo’orea NW and Tahiti NW.

*Map & temperature data*

A reef overlay map of French Polynesia was downloaded from Reefbase, where data were provided by UNEP-WCMC. The map image was manipulated in Adobe Illustrator (Figure 1A). Service d’Observation (SO) CORAIL from CRIOBE generously provided temperature data from two SBE 56 Temperature Sensors (Sea-Bird Scientific) labeled ‘B1’ and ‘P3’ at the MNW-B and MNW-F sites, respectively. Hourly temperature data at each site from June 1^st^, 2013 through June 1^st^, 2014 were extracted from their total long-term data set to correspond with the year of sampling in the current study. More information on this data set is available at SO CORAIL’s website (<http://observatoire.criobe.pf/wiki/tiki-index.php?page=Moorea>). Temperature (˚C) at each site was averaged by the time of day. In addition, the daily temperature range at each site was calculated and plotted (Figure 1B), log-transformed to meet assumptions of normality, and a one-way ANOVA was conducted to compare between reef zones. Daily mean temperature (˚C) was also calculated, normalized using the ordered quantile normalization from the *bestNormalize* package (3), and a one-way ANOVA was conducted to compare between reef zones. All analyses were carried out and visualized in R (2).

*DNA extraction*

    	Holobiont tissue was isolated by first incubating in DNA digest buffer (100 mM NaCl, 10 mM Tris-Cl (pH 8.0), 25 mM EDTA (pH 8.0), 0.5% SDS, 0.1 mg ml^−1^ Proteinase K, and 1 μg ml^−1^ RNaseA) for 1 h at 42°C, followed by a standard phenol–chloroform extraction (4). DNA concentration of each sample was quantified using a NanoDrop 1000 Spectrophotometer (Thermo Scientific) and all samples were visualized on 1% agarose gels to ensure DNA integrity. Using nuclease-free water, DNA concentrations were standardized to 25 ng/µl for 2b-RAD genotyping or to 10 ng/µl for metabarcoding of the 16S and ITS2 regions (see below).

*Coral host 2b-RAD genotyping*

    	A complete protocol for 2b-RAD genotyping, modified from (5), can be found at <https://github.com/z0on/2bRAD_denovo/>. Briefly, 100 ng of genomic DNA was digested into 36 bp fragments with restriction enzyme *BcgI* (New England Biolabs) and each fragment was multiplexed using unique 5’ and 3’ combinations of custom barcoded ligation adapters. Samples were pooled relative to gel brightness and then purified via gel extraction from a 2% agarose gel. All samples from Tahiti, along with half of Mo’orea samples were sequenced at the University of Texas at Austin on the Illumina HiSeq 2500 (76 samples total; Table S2) and libraries were spread across three lanes to increase read coverage. The remaining Mo’orea samples were sequenced on the Illumina HiSeq 4000 (60 samples total; Table S2) on 1 lane. Of these 136 samples, 10 were technical replicates (Table S2). Raw and processed read count data are available in Table S2.

*Analysis of coral host 2b-RAD data*

Bioinformatic analyses were conducted following guidelines described here: <https://github.com/z0on/2bRAD_denovo>, with modifications available at [https://github.com/nicfall/moorea_holobiont](https://github.com/NicolaKriefall/2brad_moorea).  First, all sequencing data were demultiplexed by in-line ligation barcode and adapter and barcode sequences were trimmed. Sequences were de-duplicated to remove PCR duplicates, with the exception of 21 samples (see Table S2) whose ligation adapters did not contain degenerate bases and therefore were not de-duplicated. Next, FASTX-Toolkit’s *fastq_quality_filter* tool (6) was used to filter reads containing bases with a Phred score below 20. Reads were then mapped to the *Acropora millepora* genome (7) using *Bowtie2* (8).

Analysis of Next Generation Sequencing Data (*ANGSD*) (9) was used to calculate genotype likelihoods using program defaults with the addition of the following filters: only biallelic calls, a maximum of only one best hit per read, minimum mapping quality score of 20, minimum base quality score of 25, maximum strand bias *p*-value of 10^-5^, maximum heterozygosity bias *p*-value of 10^-5^, a minor allele frequency of 0.1, a minimum depth per site of 8 reads, and sites were required to be present in at least 80% of samples. Using these settings, pairwise identity-by-state (IBS) matrices were calculated between samples, the results of which were plotted as a distance dendrogram using function *hclust()* in R (Figure S1). Technical replicates and clones identified on the dendrogram were removed, leaving only one genotype representative in the subsequent analyses. In addition, any samples with an average per-site depth of less than 7 reads were removed to improve genotype likelihood confidence (Table S2). The program *PCAngsd* (10) was used to create a covariance matrix between samples, a principal components analysis was conducted on this matrix, and the significance of multivariate dispersion and location were assessed with *vegan* functions *betadisper* and *adonis,* respectively (11). The VCF file created by *ANGSD* was subsequently converted into a BED format using *PLINK* (12) and run through *ADMIXTURE* (13) with K values 1 - 5 where the lowest cross-validation error was determined to be the optimal ‘K’. The VCF file was read into R and the package *hierfstat* (14) was used to calculate global and pairwise F_ST_ between all sites and reef zones. Finally, *BayeScan* (15) and *OutFLANK* (16) were used to identify single nucleotide polymorphism (SNP) F_ST_ outliers.

*Symbiodiniaceae metabarcoding*

From a subset (*n* = 96; 16 per reef zone per site; Table S1) of the same DNA extractions used in host genotyping, approximately 300 bp of the ITS2 region was targeted using forward primer *ITS-DINO* (5’ - TCGTCGGCAGCGTC AGATGTGTATAAGAGACAG GTGAATTGCAGAACTCCGTG - 3’), and reverse primer *ITS2Rev2* (5’ - GTCTCGTGGGCTCGG AGATGTGTATAAGAGACAG CCTCCGCTTACTTATATGCTT 3’), where underlined bases denote adapter linker, bold bases are ITS2 primer sequences (17), and the middle bases are spacer sequence. 50 ng of template DNA was added to a 15 µl PCR reaction consisting of 0.33 µM forward primer, 0.33 µM reverse primer, 0.2 mM dNTP, 1x *ExTaq* buffer (Takara), 0.025 U *ExTaq* Polymerase (Takara), 0.0125 U *Pfu* Polymerase (Agilent Technologies), and nuclease-free water. The PCR reaction profile began at 95˚C for 5 min, cycled at 95˚C for 40 s, 59˚C for 120 s, and 72˚C for 60 s for 19-24 cycles, and finished at 72˚C for 10 min. Each PCR reaction was then purified using the GeneJET PCR Purification kit (ThermoFisher) with a final elution volume of 25 µl. Each sample’s eluted PCR product was uniquely barcoded using forward (5’ AATGATACGGCGACCAC CGAGATCTACAC *NNNNNN* TCGTCGGCAGCGTC 3’) and reverse (5’ CAAGCAGAAGACGGCATAC GAGAT *NNNNNN* GTCTCGTGGGCTCGG 3’) primers, where underlined bases denote MiSeq adapter, italicized bases represent six base pair (bp) barcodes, and bold bases are adapter linker. Primers (0.33 µM) were combined with 1.5 µl of eluted product and subjected to the same PCR conditions described above, except total reaction volume was 20 µl and only 5 cycles were completed. All samples were run on a 1% agarose gel, visually assessed for relative concentrations, and pooled in corresponding concentrations. This pooled library was ethanol precipitated for purification and re-suspended in 25 µl of Milli-Q water (Millipore). 10 µl of the cleaned library was run on a 1% agarose gel containing SYBR Green (Invitrogen) and the target band was excised using a sterilized razor blade and incubated in 25 µl of Milli-Q water (Millipore) overnight at 4˚C. Liquid product was collected and submitted for 250 bp paired end sequencing on an Illumina Miseq at the University of Texas at Austin.

*Symbiodiniaceae community analyses*

    	For pre-processing, the tool *bbduk.sh* within the *bbmap* package (18) was used to remove misaligned reads containing Illumina adapter sequence and kept only reads that began with the correct ITS2 primer sequences. The program *cutadapt* (19) was then used to remove primer sequences. These pre-processed FASTQ files were read into the R environment and ITS2 analysis followed the *DADA2* pipeline (20). Reads were truncated at 220 bp and 200 bp for forward and reverse reads, respectively (based on read quality profiles), or when a quality score of less than 3 was reached. In addition, only 1 expected error was accepted, reads were required to have a minimum length of 50 bp, and reads matching the PhiX genome were removed (20). Error rates were then calculated, reads were de-duplicated, sequence variants were inferred, forward and reverse reads were merged, and bimeras were removed, following default settings (20). One sample was removed as it had 0 reads left after quality filtering (Table S1). Each of the 118 non-bimeric amplicon sequence variants (ASVs) in the resulting table was assigned taxonomy using *dada2* (20) and the ITS2 GeoSymbio database (21) with a minimum bootstrap confidence level of 70.

Next, to better evaluate the breadth of taxa represented, the LULU curation algorithm (22) was used to combine the counts of ASVs with 99% sequence matching (some ITS2 reference sequences in the GeoSymbio database differ by only a few bases) and 95% minimum relative co-occurrence. The clustered ASV table was rarefied down to 1,994 counts per sample, using *vegan* (11), to allow retaining ~90% of samples (Table S2). *Phyloseq* (23) then calculated three alpha diversity metrics: ASV richness, Shannon index, and the inverse of Simpson’s index. As these data did not meet assumptions of sampling from a normal distribution, non-parametric Mann-Whitney tests were used to assess significance between reef zones within each site. Linear regressions were used to assess correlations between alpha diversity metrics and colony size.

Next, the package *MCMC.OTU* (24) trimmed ASVs representing less than 0.1% of total reads or were present in only one sample. Counts in this final ASV table were summed by reef zone and site in a bar plot using *phyloseq* (23). In addition, a Principal Components Analysis (PCoA) using Bray-Curtis dissimilarity was conducted using *phyloseq* (23). Functions *adonis* and *betadisper* in the package *vegan* (11) were used to evaluate sample dissimilarity in multivariate space in terms of location and dispersion, respectively. These functions were each applied with 99 permutations, contrasting reef zones and colony size within each site. The function *pairwise.adonis* (from <https://github.com/Jtrachsel/funfuns>), with 99 permutations, contrasted communities between the three sites. All analyses were repeated without rarefaction, utilizing relative abundance instead to confirm results. In relative abundance analyses, all samples were retained except for three samples whose counts were at least 2.5 standard deviations lower than the mean as determined by the package *MCMC.OTU*’s *purgeOutliers* function (24) (Table S2)*.*

*Bacterial microbiome metabarcoding*

The same samples used for Symbiodinaceae ITS2 metabarcoding were used to characterize the coral’s microbiome using 16S metabarcoding, except three samples which were substituted due to insufficient remaining sample volume, and excluding three samples that failed to amplify (*n* = 93 total, 15-16 per reef zone per site; Table S1, Table S2). The bacterial 16S rRNA gene V4/V5 region was amplified using the modified forward primer Hyb515F (5’ –  TCGTCGGCAGCGTC AGATGTGTATAAGAGACAG *NNNN* GTGYCAGCMGCCGCGGTAA – 3′) and Hyb806R (5’ – GTCTCGTGGGCTCGG AGATGTGTATAAGAGACAG *NNNN* GGACTACNVGGGTWTCTAAT – 3’), where adapter linker sequence is underlined, degenerate bases are italicized, and primer sequence is in bold with degenerate bases to capture more diversity . 0.3 µM of each of these primers were combined with 20 ng of genomic DNA, 0.025 U *ExTaq* polymerase (Takara), 1x *ExTaq* buffer (Takara), 0.2 mM dNTPs, and nuclease free water up to 20 µl final volume. The PCR reaction profile cycled at 95˚C for 40 s, 58˚C for 120 s, and 72˚C for 60 s for 30 cycles, and finished at 72˚C for 10 min. Each PCR reaction was then purified using the GeneJET PCR Purification kit (ThermoFisher) and eluted to a final volume of 30 µl. Each sample’s eluted PCR product was uniquely barcoded using the same primers as in ITS2 metabarcoding (see above). Primers (0.33 µM) were combined with 1.5 µl of the eluted product and subjected to the same PCR conditions as for original 16S amplification, except 6 cycles were completed. All samples were run on a 1% agarose gel, visually assessed for relative concentrations, and 3, 5, or 10 µl of each sample was pooled accordingly. 50 µl of the pooled library was cleaned with the GeneJET PCR Purification kit (ThermoFisher) and eluted in 50 µl of elution buffer. 20 µl of the cleaned library was run on a 1.5% agarose gel containing SYBR Green (Invitrogen). The target band was excised using a sterilized razor blade and incubated in 40 µl of Milli-Q water (Millipore) overnight at 4˚C. Liquid product was collected and submitted to North Carolina State University’s Genomic Sciences Library for 250 bp PE sequencing on the Illumina MiSeq.

*Bacterial microbiome analyses*

    	Pre-processing using the tool *bbduk.sh* within the *bbmap* package (18) was used in the same manner as for ITS2 analyses, with an additional step of removing four bases at the beginning of each read introduced by degenerate primers. Analysis of pre-processed FASTQ files was conducted in R following the *dada2* pipeline (20) in the same manner as for ITS2, except that reads were truncated at 200 and 180 bp for forward and reverse reads, respectively. Taxonomy was assigned using default settings and the Silva version 132 dataset formatted for *dada2* (20,25).

Using *phyloseq* (23), ASVs that were assigned to family “Mitochondria”, order “Chloroplast”, or did not assign to kingdom “Bacteria” were removed. Using *vegan* (11)*,* the ASV table was rarefied down to 12,000 reads, as this value would permit retaining ~90% of the total samples (*i.e.* 9 samples were removed for having fewer reads; Table S2). *Phyloseq* (23) was used to calculate three diversity metrics: ASV richness, Shannon index, and the inverse of Simpson’s index. ASV richness and inverse Simpson index were log-transformed to meet assumptions of normality and then a one-way ANOVA with Tukey HSD post hoc test was conducted to compare all diversity metrics across sites and reef zones. In addition, linear regressions assessed correlations between alpha diversity metrics and colony size. *MCMC.OTU* (24) then trimmed ASVs that represented less than 0.01% of total counts or were present in only one sample. This final ASV table was summarized in a bar plot using *phyloseq* (23).

Using the final ASV table, *adonis* and *betadisper* functions in the package *vegan* (11) were used to evaluate sample dissimilarity in multivariate space in terms of location and dispersion, respectively. These functions were applied comparing reef zones and colony size within each site. Then, total microbiome results were separated into core (present in >70% of samples) and accessory microbiomes using package *microbiome* (26). *Adonis* and *betadisper* analyses were repeated comparing reef zones and colony size within each site for the core and accessory microbiomes. Analyses were again repeated using unrarefied, relative ASV abundance per sample to confirm all results.

Next, the package *indicspecies* (27) was used to find significant associations between ASVs and reef zones at each site with 999 permutations, followed by a Benjamini & Hochberg multiple test correction (28). Finally, the online resource *Piphillin* (29) predicted the metagenomic content of the samples using the Kyoto Encyclopedia of Genes and Genomes (KEGG) Database (30). *DESeq2* (31) was used to identify differentially enriched metagenomic content across reef zones within sites.

**RESULTS**

*Temperature data*

Daily temperature range was significantly higher in back reef (0.93 ± 0.45 ˚C; mean ± SD) relative to fore reef (0.51 ± 0.23 ˚C; mean ± SD) environments at Mo’orea NW (Figure 1B; one-way ANOVA: F_1,728_ = 306.2, *p* < 0.001). In contrast, mean daily temperature (˚C) did not vary between reef zones.

*Coral colony surface area*

Coral colony area (cm^2^) was significantly higher at back reef locations relative to fore reef at both Moorea NW (one-way ANOVA: F_1,48_ = 22.81, *p* < 0.001) and Tahiti NW (one-way ANOVA: F_1,44_ = 11.82, *p* < 0.01) (Figure 1C).

*Symbiodiniaceae community composition*

Within all three sites, fore reef and back reef communities were significantly different (PERMANOVA: Mo’orea NW: F_1,25_ = 2.18, *p* < 0.05; Mo’orea SE: F_1,29_ = 6.35, *p* < 0.05; Tahiti NW: F_1,23_ = 3.1, *p* < 0.05) (Figure 2A). Data were not significantly dispersed in multivariate space, with the exception of Tahiti NW, where back reef data were more dispersed than fore reef data (PERMDISP: F_1,25_ = 7.64, *p* < 0.05) (Figure 2A). Colony size significantly influenced community data at Tahiti NW (PERMANOVA: F_1,23_ = 4.06, *p* < 0.05), but not at Mo’orea NW, and there were no significant interactions between colony size and reef zone at either site. With reef zone data combined by site, algal communities were different between Mo’orea NW and Tahiti NW (PERMANOVA: F = 5.74, *p* < 0.01), and between Mo’orea SE and Tahiti NW (PERMANOVA: F = 13.67, *p* < 0.01), but not between Mo’orea SE and Mo’orea NW (Figure S4B).

Two metrics of alpha diversity indicated higher overall diversity in fore reef algal symbionts when compared to back reef (Wilcoxon rank sum tests: Shannon: W = 572, *p* < 0.01; Simpson: W = 584, *p* < 0.01). Within each site, there was higher diversity in fore reef samples at Mo’orea NW (Wilcoxon rank sum tests: Shannon: W = 50, *p* = 0.053; Simpson: W = 50, *p* < 0.05) and Mo’orea SE (Wilcoxon rank sum tests: Shannon: W = 47, p < 0.01; Simpson: W = 57, *p* < 0.05), but no differences were observed across reef zones at Tahiti NW (Figure 2C). There was only a significant difference in ASV richness across reef zones at Mo’orea SE, with higher ASV richness in the fore reef (Wilcoxon rank sum test: W = 60, *p* < 0.05). Diversity metrics were not significantly correlated with colony size.

*Bacterial community composition*

Multivariate location of 16S communities was significantly different between Mo’orea SE and Mo’orea NW (PERMANOVA: F = 2.91, *p* < 0.05) and between Mo’orea SE and Tahiti NW (PERMANOVA: F = 3.45, *p* < 0.01), but not between Mo’orea NW and Tahiti NW (Figure S7).

Within each site, 16S communities were significantly different between the fore reef and back reef at Mo’orea SE (PERMANOVA: F_1,28_ = 3.975, *p* < 0.001) and Tahiti NW (PERMANOVA: F_1,25_ = 4.32, *p* < 0.001), but not at Mo’orea NW (Figure 3A). There were no significant differences in multivariate dispersion across reef zones or sites. Patterns did not change when examining the core microbiome (Table S4). However, all reef zone comparisons at all three sites were significant when examining the accessory microbiome (PERMANOVAs: Mo’orea NW: F_1,25_ = 1.86, *p* < 0.05; Mo’orea SE: F_1,28_ = 2.78, *p* < 0.001; Tahiti NW: F_1,25_ = 2.72, *p* < 0.001). There were no significant relationships between colony size and community composition and no interactions between colony size and reef zone.

**Table S1.** Site information: sample numbers sequenced followed by numbers retained after quality control in parentheses for 2b-RAD (coral host), ITS2 (algal symbiont), and 16S (bacterial).

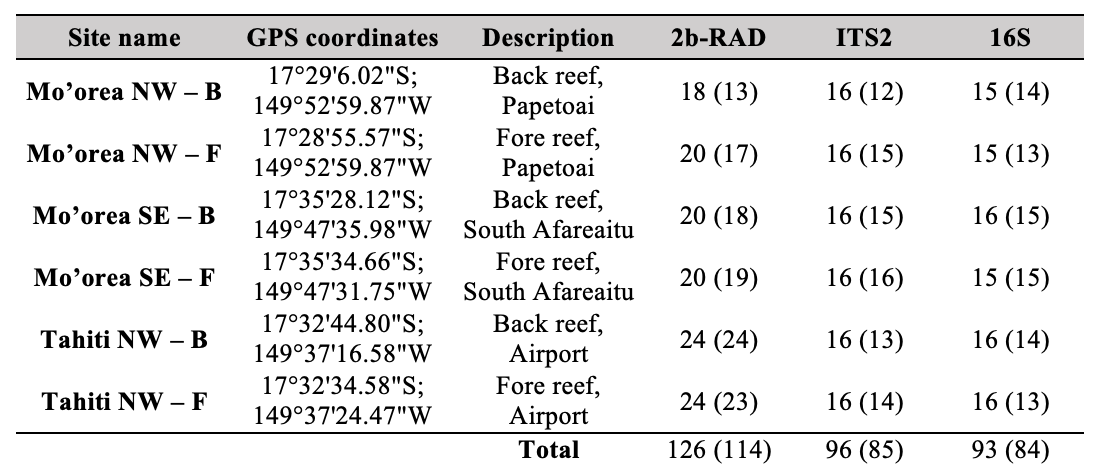

**Figure S1.** Identity-by-state (IBS) pairwise comparison results from all samples. Sample names followed by a ‘d’ indicate a technical replicate. In addition, samples MNW-F_125 and MNW-B_58 are strongly clustered with between-sample similarities on par with technical replicates, indicating that they are incidental clones. Technical replicates and clones, which all clustered below the 0.25 dendrogram height threshold (dashed red line), were removed from downstream analyses and are indicated in red.

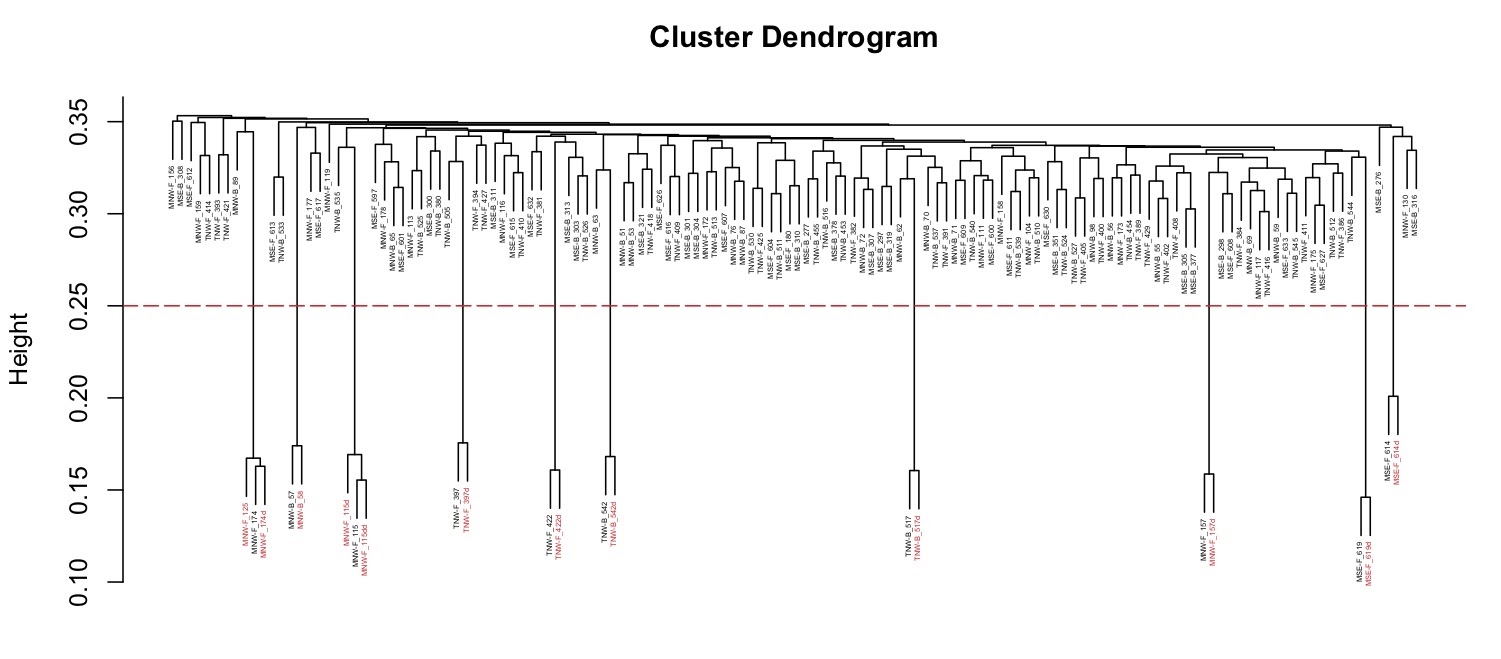

**Figure S2.** Principal components (PC) analysis of coral host 2b-RAD data, displaying overlap of samples across **(a)** reef zones (BR: back reef, FR: fore reef) and **(b)** sites (MNW: Mo’orea NW, MSE: Mo’orea SE, TNW: Tahiti NW).

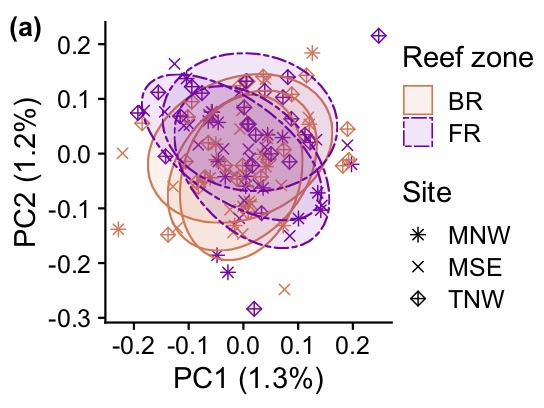

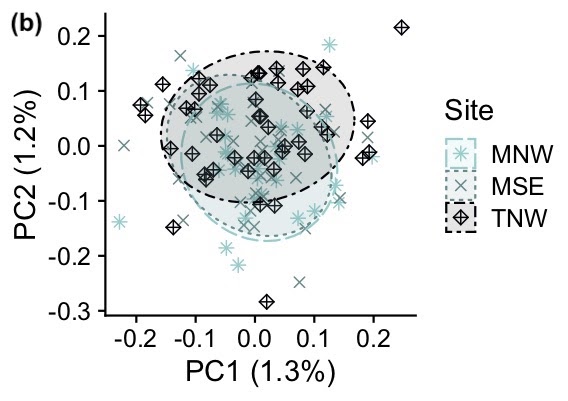

**Figure S3.** Genetic structure across sites (MNW: Mo’orea NW, MSE: Mo’orea SE, TNW: Tahiti NW) and reef zones (F: fore reef, B: back reef) from *ADMIXTURE* ancestry estimation with K = 2 (optimal K = 1). Individuals are plotted on the x-axis, with assignment probability to one of two ancestral groups plotted on the y-axis.

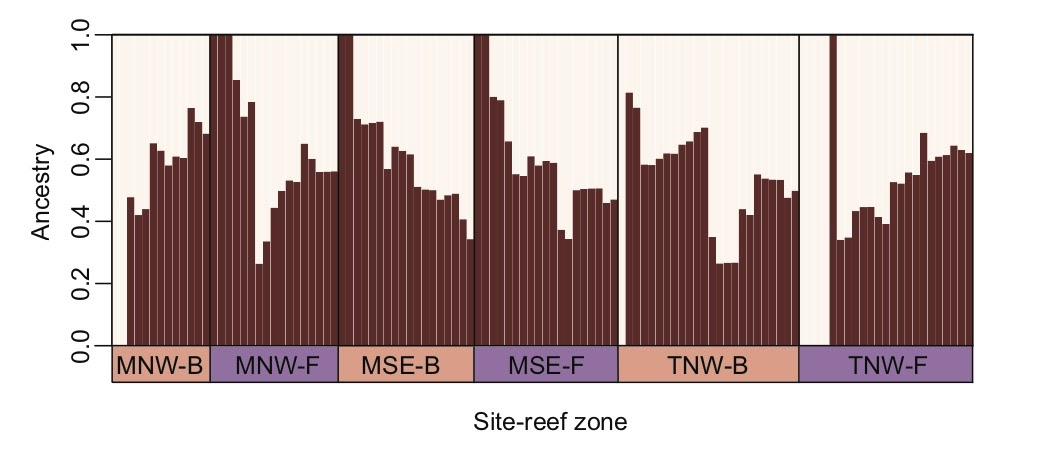

**Table S3.** Pairwise F_ST_ comparisons between all sites (MNW: Mo’orea NW, MSE: Mo’orea SE, TNW: Tahiti NW) and reef zones (B: back reef, F: fore reef).

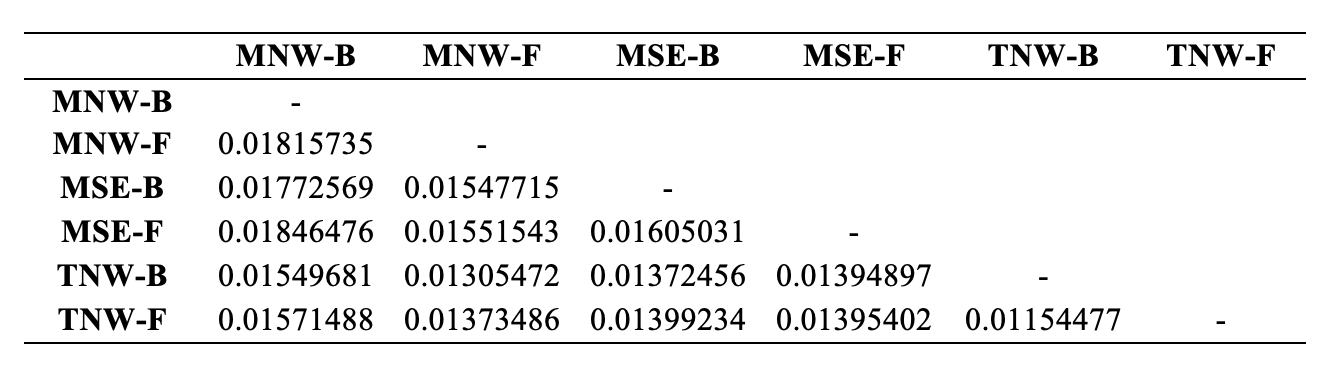

**Figure S4. (a)** Relative symbiont abundance summed by reef zone (BR: back reef, FR: fore reef) and site. ITS2 types are given in plot legend, where more than one ASV assigned to the same ITS2 type are numbered (“C.” represents ASVs identified as  *Cladocopium* of unknown type). **(b)** Multivariate ordination (PCoA) based on Bray-Curtis dissimilarity between samples. Unique colors and shapes represent Symbiodiniaceae communities at each site (MNW: Mo’orea NW, MSE: Mo’orea SE, TNW: Tahiti NW). Significance of pairwise comparisons (*pairwise.adonis*) between sites are provided in the top left.

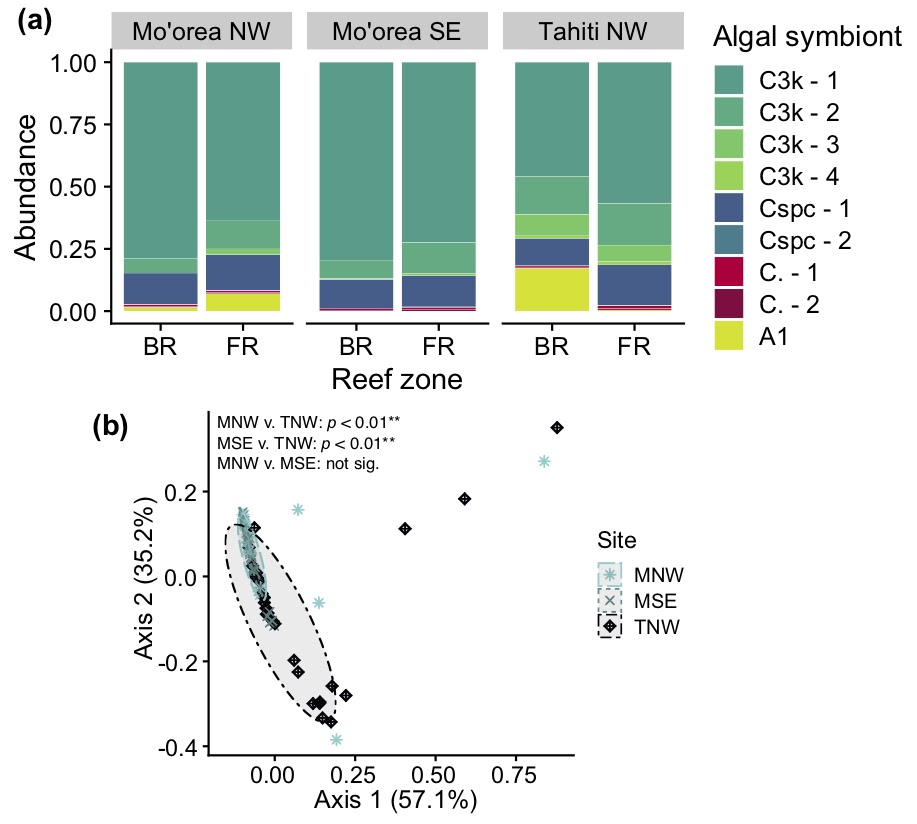

**Figure S5.** Total counts of Symbiodiniaceae ASVs across reef zones (BR: back reef, FR: fore reef) in French Polynesia at **(a)** Mo’orea NW **(b)** Mo’orea SE, and **(c)** Tahiti NW. ITS2 types are indicated in individual panel titles, where more than one ASV assigned to the same ITS2 type are numbered and “C.” represents ASVs assigned *Cladocopium* but unassigned at the type level.

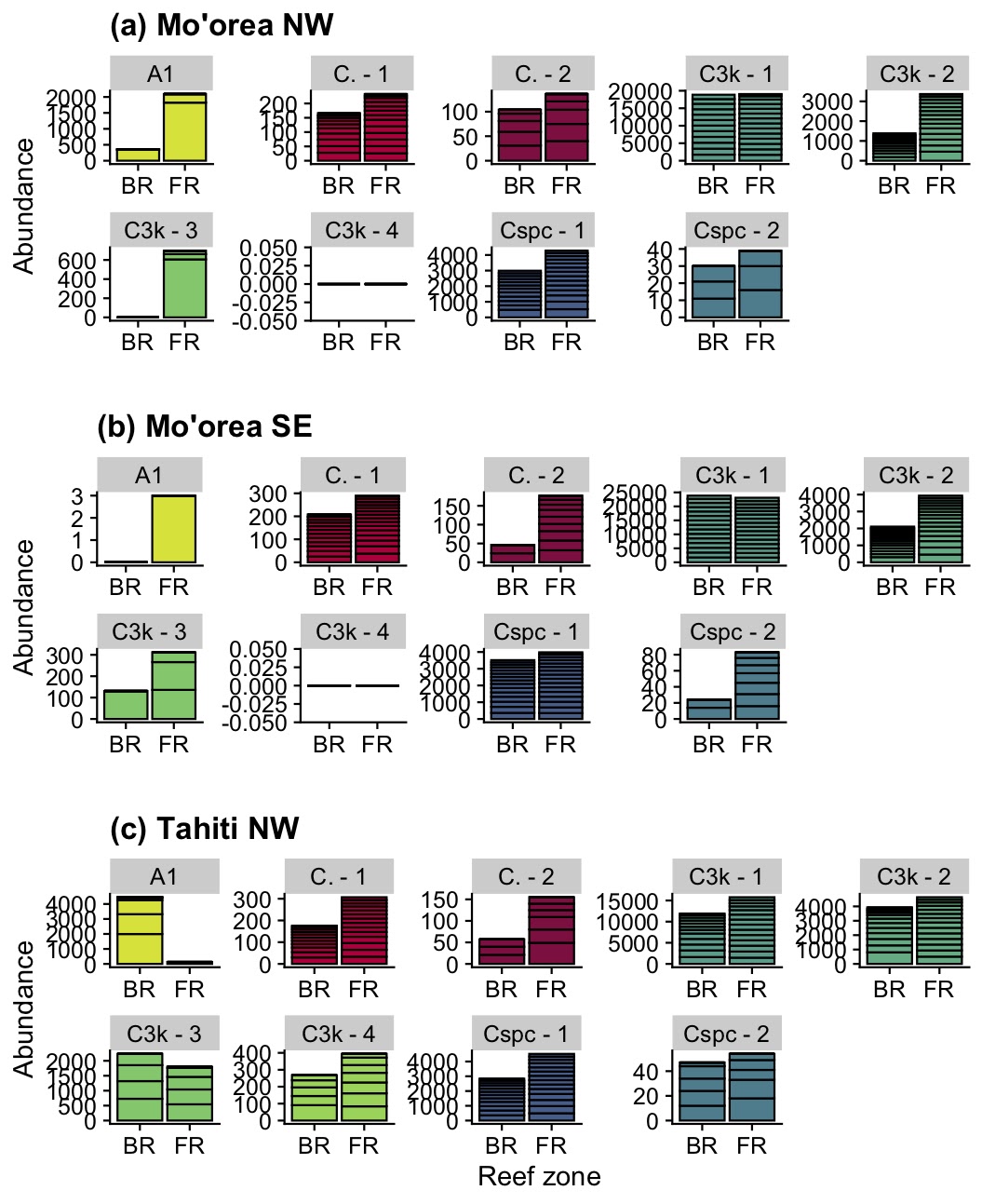

**Figure S6.** Bacterial community abundances, summed across reef zones (BR: back reef, FR: fore reef) and sites (panel titles), colored by bacterial Class.

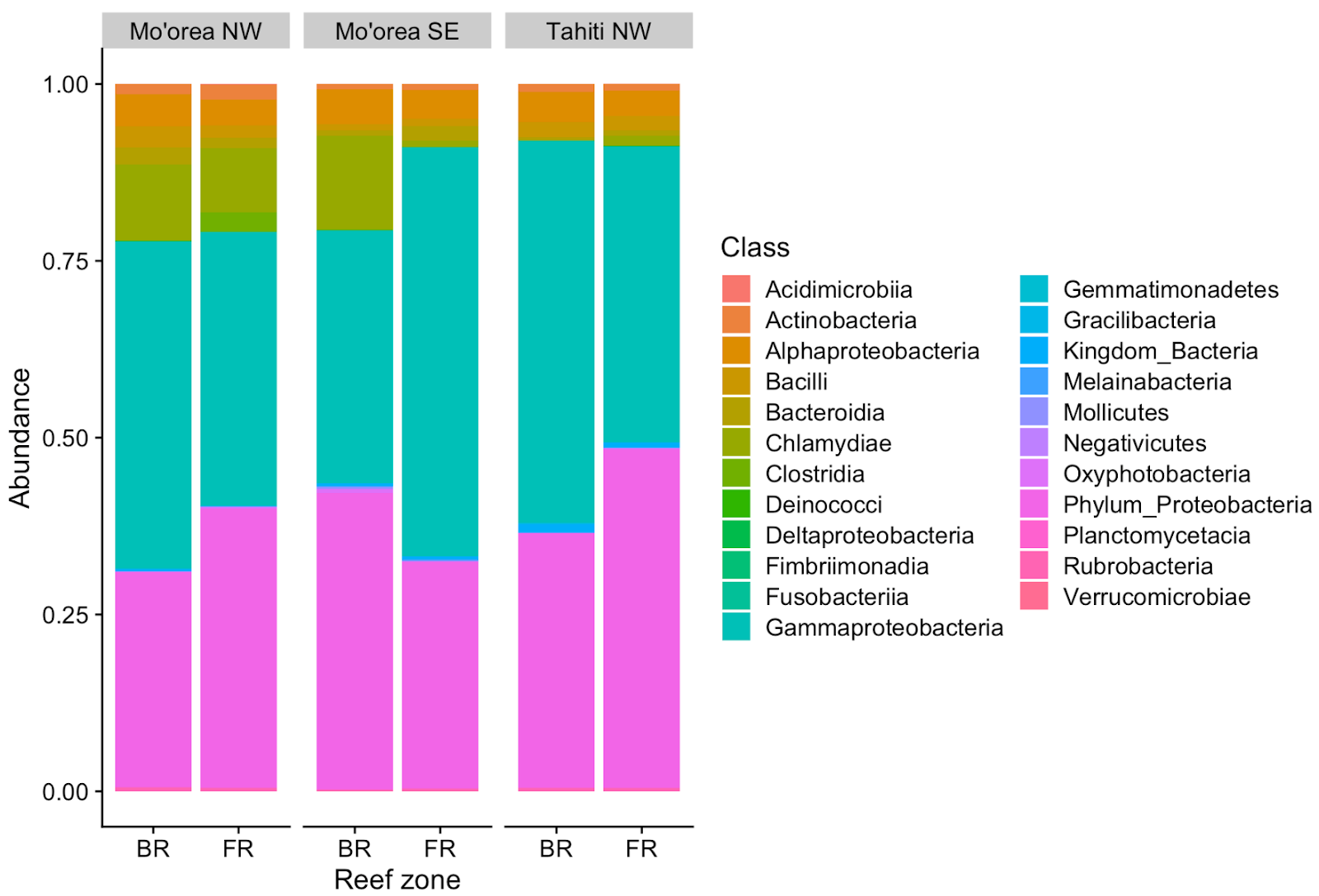

**Table S4.** Identity information of core (present in >70% of samples) bacterial members of *Acropora hyacinthus* and their average relative abundance across all samples.

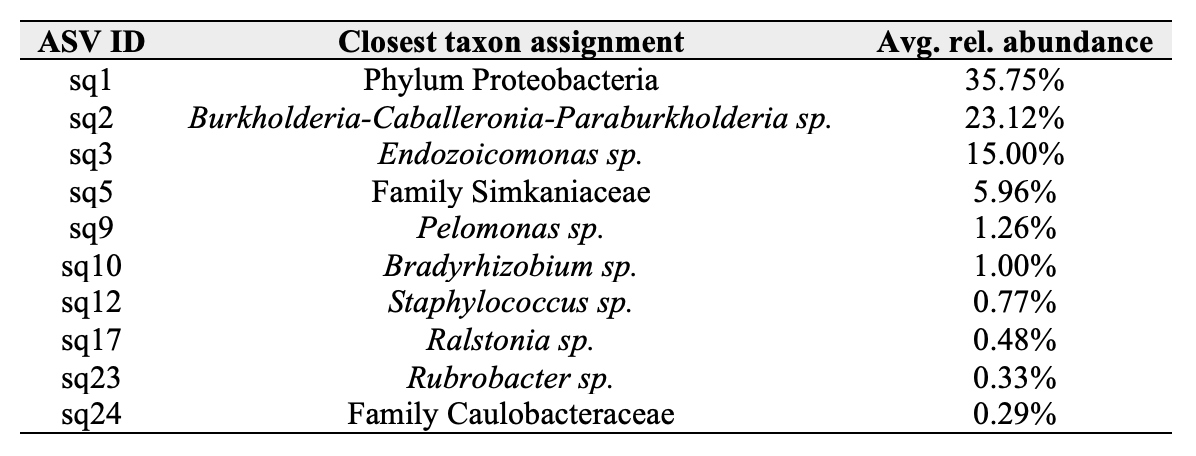

**Figure S7.** Bacterial communities in *Acropora hyacinthus* across sites (MNW: Mo’orea NW, MSE: Mo’orea SE, TNW: Tahiti NW). This plot shows multivariate ordination (PCoA) based on Bray-Curtis dissimilarity between samples. Significance of pairwise comparisons (*pairwise.adonis*) between sites are provided in the top right.

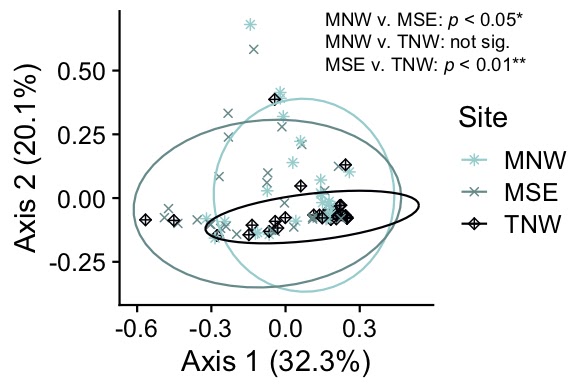

**Table S5.** KEGG functions enriched in reef zones (B: back reef, F: fore reef) and shared across more than one site (MNW: Mo’orea NW, MSE: Mo’orea SE, TNW: Tahiti NW).

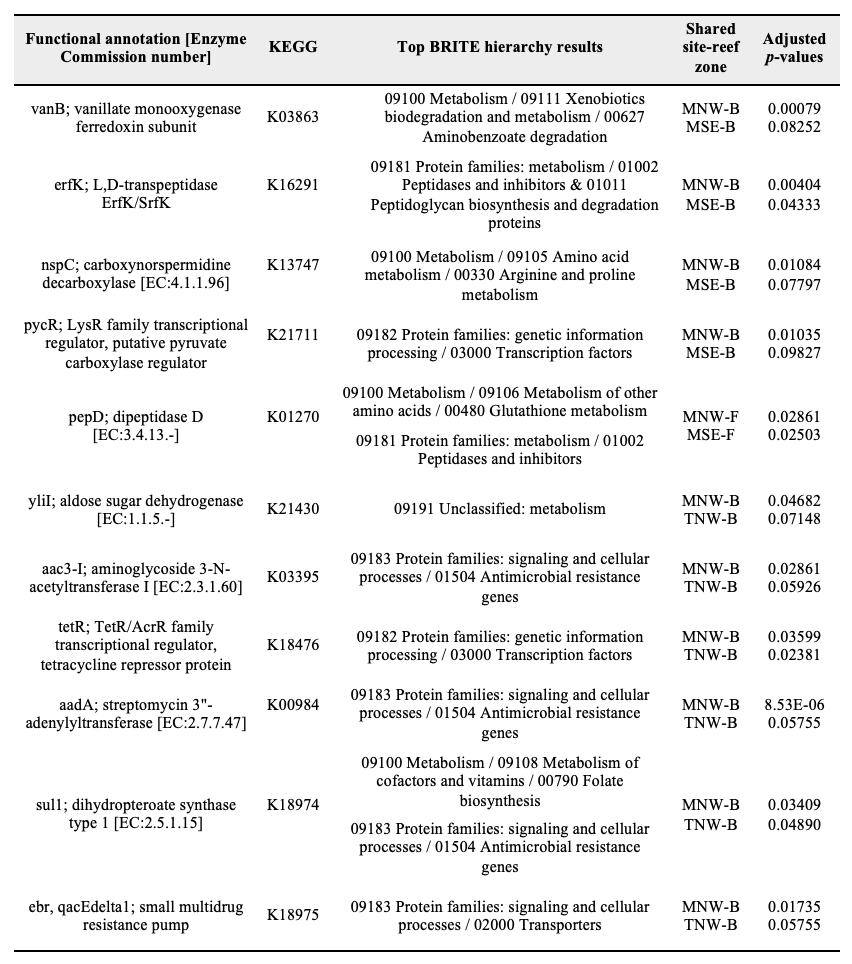
